# RiboFlow v2: a configurable ribosome profiling pipeline reveals measurements sensitive to read-mapping methodology

**DOI:** 10.64898/2026.10.05.756038

**Authors:** Daniel Nguyen, Can Cenik

## Abstract

Ribosome profiling experiments capture a snapshot of translation by sequencing ribosome-protected fragments (RPFs). Alignment to a well-curated transcriptome captures most RPF signal, but can miss ribosome activity when expressed exons are absent from the reference. Genome alignment of RPFs expands the detectable sequence space, but can expose ambiguity between homologous loci. Existing workflows often couple reference choice with the handling of ambiguously mapped reads, making it difficult to isolate the effects of either decision on downstream analyses. Here, we developed RiboFlow v2 by adding genome alignment and configurable handling of multimapping reads to the existing transcriptome workflow. Using 24 human ribosome profiling libraries and their matched RNA-seq data, we found that nucleotide-level coverage at matched coding-sequence positions and translation efficiency (TE) were broadly concordant between alignment strategies under stringent alignment filters. Even under these conditions, individual genes showed substantial differences in coverage and TE estimates. These differences included coverage in alternative exons absent from selected transcripts and ambiguous alignments between protein-coding genes and processed pseudogenes. Clustering genes by read-assignment patterns identified groups associated with pseudogene prevalence and highlighted shared exonic sequence among reference transcripts as a source of alignment discrepancies. RiboFlow v2 extends the analysis beyond selected reference transcripts and enables identification of genes whose ribosome coverage and abundance estimates depend on reference composition and how ambiguously mapped reads are handled.

## 1. INTRODUCTION

Ribosomes can shield pieces of mRNA from RNase digestion, and these ribosome-protected fragments (RPFs) can be sequenced to capture a snapshot of which transcript regions are being translated [1], also known as translons [2]. The sequenced RPFs are aligned to a reference transcriptome or genome [3,4]. From these alignments, A/P-site offsets are inferred [5] and applied to RPFs for downstream analyses such as ribosome stalling, codon occupancy, and translon detection [2], all of which depend on accurate read assignment [6]. RPF read assignment depends on the reference used and how reads that align equally well to multiple locations are handled.

Restricting alignment of ribosome profiling reads to a transcriptome assumes that selected transcripts represent the isoforms expressed in the sample. However, isoform usage varies across tissues and cell types [7,8], and isoform transcripts differ in their translational output [9]. RPFs from expressed sequences that are absent from the transcriptome reference will not be accurately assigned. In RNA-seq, such reads fail to align or are spuriously mapped [10]. For ribosome profiling, this may mean transcriptome-based pipelines will miss ribosome activity for transcripts excluded from the reference. Aligning RPFs to the genome addresses this limitation because it includes all possible isoform exons.

The genome, however, includes additional homologous loci. Protein-coding genes in the human genome can share sequence similarity with other protein-coding genes, pseudogenes, and lncRNA loci [11]. Because RPFs are shorter than RNA-seq reads, they more often align equally well to multiple loci [12]. Retaining multi-mapping reads can create artificial peaks in ribosome occupancy that resemble translational pausing [12], whereas discarding them may remove genuine ribosome occupancy. Methods exist to rescue or fractionally assign multi-mapping reads [13] but aren’t standard in ribosome profiling pipelines. Genome alignment removes the dependence on transcript selection but introduces ambiguity from multimapped RPFs.

Mapping methodology decisions may also affect translon discovery. Ribosome profiling has identified translons in previously unannotated regions [14], including ones that encode microproteins or immunogenic peptides that may serve as tumor antigens [15]. Large-scale translon discovery studies commonly align RPFs to the genome [16–19]. However, translon discovery is not limited to genome-based pipelines, and translon sets identified by transcriptome and genome pipelines can differ [20]. Regardless of the alignment reference, translon identification requires distinguishing ribosome-protected RNA fragments from those originating from other RNA-protein complexes or stable RNA secondary structures [21]. To distinguish these sources, translon identification tools typically rely on translation-specific signals such as three-nucleotide periodicity [6,9,22,23] or uniform A/P-site coverage indicative of translation elongation [24]. Because these signals are inferred from ribosome profiling read alignments, their accuracy depends on whether coverage profiles reflect true ribosomal activity. Translon discovery studies typically retain only uniquely mapped ribosome profiling reads [17– 19]. Restricting genomic alignments to uniquely mapped RPFs has been shown to impact the identification of some translons with proteomic support [25]. Together, these studies show how the handling of multi-mapping ribosome profiling reads during alignment can influence translation-specific signals and translon identification.

In addition to translon identification, ribosome profiling is most frequently used to study translational regulation. Paired with RNA-seq, many studies approximate translation efficiency (TE), estimated as the log-ratio of RPF to mRNA read counts [26]. These estimates similarly depend on accurate read assignment. In RNA-seq, restricting analyses to uniquely mapped reads underestimates the abundance of protein-coding genes relative to their pseudogenes [27], and reference choice alters transcript abundance estimates [10]. Because RPFs map to multiple competing loci more frequently [12] than RNA-seq reads, reference and multi-mapping choices may exacerbate these effects in ribosome profiling data, as TE and abundance estimates may be particularly sensitive for genes with alternative isoforms or homologous loci.

Existing ribosome profiling pipelines couple alignment reference choice and multi-mapping policy in different ways. RiboFlow and riboviz2 use transcriptome alignment and matched RNA-seq [28,29], whereas genome-based pipelines vary in how they retain multimapping genomic alignments [30,31]. Pipelines such as nf-core/ribo-seq [32] project genome alignments to transcript coordinates. Projection can introduce ambiguity when a single genomic alignment fits multiple transcript models. Although probabilistic transcript counting assigns fractional weights to reads compatible with multiple transcripts, nf-core/riboseq does not incorporate these weights when riboWaltz [5] analyzes projected alignments for A/P-site inference. The original RiboFlow relies on transcriptome alignment [29] and cannot evaluate RPFs from alternative isoforms or ambiguity among homologous genomic loci. Because existing pipelines do not allow independent configuration of the alignment reference and multi-mapping policy, their workflows cannot easily distinguish their separate effects on positional signals and TE estimates.

To isolate the effects of alignment reference and multi-mapping policy and examine ribosome profiling signal beyond selected transcripts, we developed RiboFlow v2, an updated version of RiboFlow in Nextflow DSL2 [33]. RiboFlow v2 retains transcriptome alignment while adding genome alignment and configurable handling of multi-mapping reads. We compared read assignments, positional RPF signals, and abundance estimates across alignment routes and mapping policies, and grouped genes by read-assignment patterns to examine their associations with pseudogenes and transcript features. Although measurements broadly agreed within selected reference CDSs, individual genes remained sensitive to alignment methodology. Comparing genome and transcriptome alignments highlighted how transcript selection and the exclusion of competing loci affect coverage and read assignment.

## 2. DESIGN AND IMPLEMENTATION

RiboFlow is a Nextflow-based pipeline for the uniform processing of ribosome profiling (Ribo-seq) data, with optional matched RNA-seq data. The original workflow aligned RPFs to a transcriptome reference and summarized them in a compact file format (.ribo), separated by read length, for use with RiboR, RiboPy [29], and RiboGraph [34]. Here we present RiboFlow v2, a port of RiboFlow from Nextflow DSL1 to DSL2 that retains the transcriptome path and adds a genome alignment path using STAR v2.7.11b [35], multithreaded unique molecular identifier (UMI) deduplication with UMICollapse [36], and optional strand-separated coverage tracks (bigWig). All validation analyses below were produced using the genome-alignment path and compared against the transcriptome-aligned .ribo files.

### 2.1 Pipeline overview

RiboFlow v2 is implemented in Nextflow DSL2 (v26.04.2; Java 23.0.2) using composable subworkflows and per-process modules. A single YAML parameter file specifies whether to run genome alignment, transcriptome alignment, or both, and independently enables optional matched RNA-seq processing. Preprocessing is shared across the selected alignment routes and includes adapter trimming, optional UMI extraction, and contaminant filtering. Preprocessed reads are then aligned, quality-filtered, merged within each sample group, deduplicated, and converted into route-specific analysis outputs (Fig 1).

**Fig 1.**
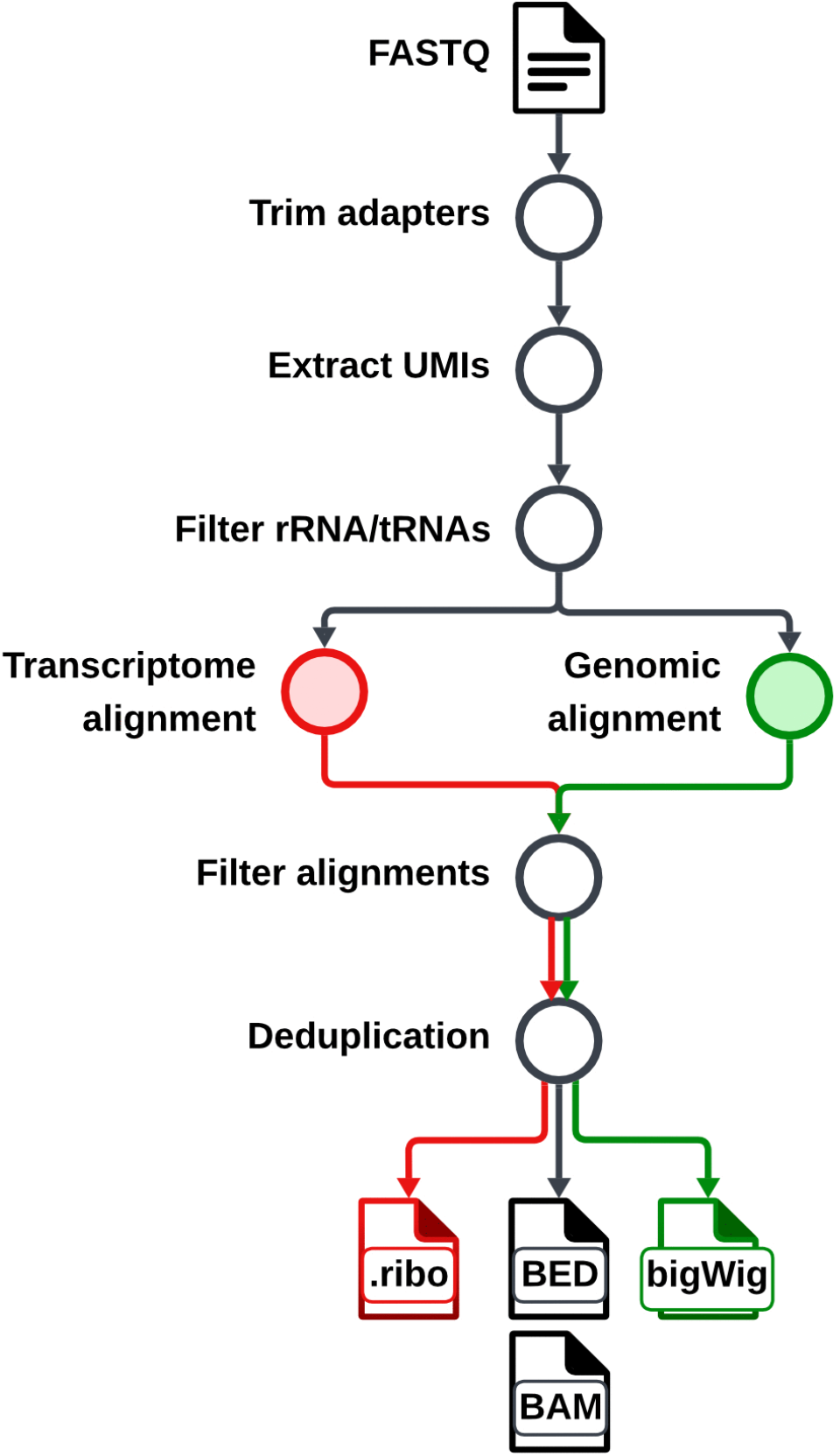
Overview of the RiboFlow v2 pipeline. Ribosome profiling reads, with optional matched RNA-seq, are adapter- and quality-trimmed, UMI-extracted, and filtered against an rRNA/tRNA contaminant index. Surviving reads are aligned in parallel to the genome (green) and the transcriptome (red). Alignments from both routes are quality-filtered and deduplicated, producing indexed BAM and BED files. The transcriptome route produces .ribo files, and the genome route produces optional bigWig files.

### 2.2 Inputs and reference data

Ribosome profiling inputs are one or more single-end FASTQ files per sample group. Matched RNA-seq inputs are optional and may comprise one or more single- or paired-end FASTQ files per sample group. Pre-alignment contaminant removal requires an rRNA/tRNA FASTA and corresponding Bowtie2 [37] index. For this study, we reused the human v2 rRNA/tRNA filter file generated from [29].

Both the transcriptome and genome-alignment references were based on the GRCh38.p13 human genome assembly and GENCODE v34 annotation. For transcriptome alignment, we reused the RiboFlow human v2 transcriptome reference, generated with a custom Python script (generate_appris_reference.py) [29]. The reference contains protein-coding APPRIS principal transcripts with valid start and stop codons. When multiple transcripts for a gene were classified as APPRIS principal isoforms, the longest transcript was retained. Of 27,315 candidate transcripts, 7,579 were removed through transcript filtering, resulting in a reference transcriptome containing 19,736 transcripts. The same script also produced the annotation inputs required for .ribo file generation, enabling quantification of coverage by transcript regions and read lengths.

Genome alignment requires a STAR index, which may be supplied prebuilt or generated within the workflow using parameters specified in the configuration YAML file. For this study, the index was generated using --sjdbOverhang 28.

### 2.3 Read preprocessing and contaminant removal

Read quality can optionally be assessed with FASTQC [38] v0.12.1. Cutadapt [39] v5.2 then trims adapters and low-quality bases using arguments specified in the YAML parameter file. When UMI-based deduplication is enabled, UMI-tools [40] v1.1.6 extracts the UMI and appends it to the read header. Trimmed reads are aligned with Bowtie2 against the rRNA/tRNA contaminant index, and only unaligned reads are passed to the selected alignment paths.

### 2.4 Genome alignment with STAR

Reads that pass contaminant filtering are aligned to the genome with STAR, using parameters set separately for ribosome profiling and RNA-seq. SAMtools [41] v1.23 optionally applies a mapping-quality filter. Filtered alignments are merged within each sample group and processed by the selected deduplication method. SAMtools produces a sorted, indexed BAM, and bedtools [42] v2.31.1 generates a BED representation. Optionally, deepTools [43] v3.5.5 bamCoverage generates strand-separated bigWig coverage tracks from the genome alignments.

### 2.5 Transcriptome alignment and .ribo file generation

When transcriptome alignment is enabled, RiboFlow follows the previously described workflow for transcriptome alignment, filtering, deduplication, and .ribo file generation [29], reimplemented in Nextflow DSL2. For this study, transcriptome alignments were restricted to MAPQ of at least 10 before deduplication.

### 2.6 Deduplication

Deduplication is optional and is applied after merging lanes by sample. In position-based mode, RFCommands [29] collapses reads with identical alignment coordinates in the merged BED file. In UMI-based mode, UMICollapse v1.1.0 collapses reads in the merged BAM when they share both the same alignment coordinates and a UMI. The libraries analyzed in this study used position-based deduplication as they lacked UMIs.

### 2.7 Methods for Results

#### 2.7.1 Data Processing

Genomic alignments were restricted to uniquely mapped reads and transcriptome alignments to MAPQ of at least 42. Unmapped, secondary, and supplementary alignments were excluded. Read lengths were selected independently for each alignment route as described in [44], using 21-40nt CDS-assigned RPFs (qc_core.py::select_read_lengths()). P-site offsets were estimated independently for each selected read length and alignment route using psite_offset.py [45].

The selected reference transcripts defined the exon and CDS coordinates used for comparisons. CDSs extend from the first nucleotide of the start codon inclusive to the first nucleotide of the stop codon (exclusive).

#### 2.7.2 Coverage profile construction and comparison

For each library and alignment route, we constructed P-site and whole-RPF coverage profiles from RPFs in the selected read-length window. For P-site coverage, assigned RPFs contributed one count at their inferred P-site, and whole-RPF coverage contributed counts at aligned positions overlapping exons of the selected reference transcript, without P-site offsetting. Genomic alignments were projected to 5’-to-3’ spliced transcript coordinates by concatenating exons from selected transcripts (transcript_coords.py). Transcriptome alignments already used these coordinates.

Correlations between alignment routes were calculated across matched positions after excluding 15 nucleotides from each end of the CDS to reduce the influence of initiation and termination peaks [46]. Spearman correlations were calculated from raw coverage counts, and Pearson correlations were calculated from log2(count + 1)-transformed coverage. Transcript-level correlations were calculated for transcripts with coverage in both routes. Library-level correlations were calculated after concatenating coverage across all matched interior CDS positions, including positions with zero coverage in both routes.

#### 2.7.3 Translation efficiency estimation and comparison

For each sample and alignment route, RPFs in selected read-length windows and RNA-seq reads of any length were counted when their 5’ ends aligned to a selected transcript CDS. Genome-aligned reads whose 5′ ends fell within the CDSs of more than one selected transcript were excluded as ambiguous.

Counts per million (CPM) were calculated separately for each assay and alignment route. Transcripts were retained if they exceeded 1 CPM in both RPF and RNA-seq data under both alignment routes in at least half of the cell lines. This filtering retained 11,589 of the 19,736 reference transcripts. For each assay, median-of-ratios normalization factors were calculated per cell line from the geometric mean of genome- and transcriptome-aligned counts. Factors were estimated using transcripts with nonzero counts for both assays and alignment routes across cell lines.

TE was calculated in cell lines with nonzero counts as the log_2_ ratio of normalized RPF to mRNA counts [1]. ΔRNA, ΔRibo, and ΔTE were defined as genome minus transcriptome. By definition, ΔTE equals ΔRibo minus ΔRNA. Because the same normalization factor was applied to both alignment routes within each assay, these differences were equivalent to the corresponding differences in log2 counts.

For each transcript, we used a two-sided one-sample t-test to determine whether ΔTE values across eligible cell lines differed from zero. We calculated 95% confidence intervals using the t distribution with n-1 degrees of freedom and adjusted p-values using the Benjamini-Hochberg procedure [47].

#### 2.7.4 Comparison of read assignment and genomic multimapping, and gene clustering

Deduplicated ribosome profiling BAM files from aligner routes were compared by unique read IDs without read-length filtering. Each library’s read-ID union was partitioned into shared, genome-only, and transcriptome-only (TO) reads. Transcriptome-aligned reads with MAPQ of at least 10 to their selected reference transcript were considered. Shared and genome-only reads were further classified as uniquely mapped (SH-U and GO-U) or multimapped (SH-M and GO-M) to the genome.

Primary genomic alignments with secondary alignments that had tied alignment scores were compared. For each library, the proportion of SH-M reads whose tied placements had 5′ ends in a protein-coding exon and a processed-pseudogene exon was calculated. GO-U reads were classified as overlapping omitted alternative exons if at least one aligned base of their genomic alignment fell within a gene’s exonic sequence absent from a selected transcript.

For gene-level analyses, the read-ID union of a gene consisted of distinct read IDs with a highest-scoring genomic alignment at the gene or a transcriptome alignment to its selected reference transcript. Each read was counted once per gene but could belong to multiple gene unions if highest-scoring genomic alignments were tied. Gene-specific pseudogene ties additionally required one of the 5′-end alignments to fall in an exon of the gene examined.

P-site coverage of LRRFIP1 used projected alignments onto the combined exonic coordinates of the selected and alternative isoforms. Genomic P-site coverage included unique reads and multi-mapping reads using their highest-scoring alignment at the locus.

Genes in GSM2100602 with at least 100 associated read IDs were hierarchically clustered by their five read-assignment proportions using Euclidean distance and Ward’s linkage. The dendrogram was cut into four clusters, and centroids were the mean proportions of member genes. Each cluster quantified genes with at least one (processed or unprocessed) pseudogene, using the two-way consensus of the Yale and UCSC predictions for GENCODE v34.

To measure exonic sequence absent from the selected reference transcript, we merged the exons of all GENCODE v34 transcripts for each gene and calculated the percentage of bases in that union not covered by the selected transcript’s exons. To measure overlap between selected transcripts of different genes, we calculated the percentage of each selected transcript’s exonic bases whose genomic coordinates overlapped an exon of another gene’s selected transcript on the same chromosome and strand.

## 3. RESULTS

### 3.1 P-site estimation and CDS periodicity are consistent across alignment methods

We selected 24 human ribosome profiling samples from RiboBase [44], a database of curated ribosome profiling and RNA-seq samples (S1 Table). Each library was processed with RiboFlow v2 using both alignment routes (Methods). The selected read-length windows were identical between alignment routes for 22 of the 24 libraries (Fig S1A). Among these shared read lengths, the fraction of CDS P-sites in the first frame differed between alignment methods by a mean absolute value of 0.33 percentage points, with individual differences ranging from −0.4 to +2.0 percentage points (Fig S1B). Thus, the two alignment routes produced similar results for read-length selections, P-site offsets, and CDS P-site periodicity for these libraries.

### 3.2 Ribosome profiling coverage is broadly concordant across alignment strategies

Alignment routes may differ in nucleotide-level coverage patterns within individual transcripts, despite consistent overall P-site estimation and aggregate CDS periodicity. COMT represented a typical mRNA, with its P-site distribution having a Spearman ρ of 0.961 (Fig 2A, 2C) between the two alignment routes. In contrast, GAPDH is an example of an outlier gene that showed low agreement (P-site Spearman ρ = 0.091; Fig 2B and 2C). Thus, although P-site coverage was highly concordant for the typical transcript (Fig 2D), individual transcripts could show substantial differences in their coverage profiles. Such discrepancies may affect analyses based on positional coverage.

**Fig 2.**
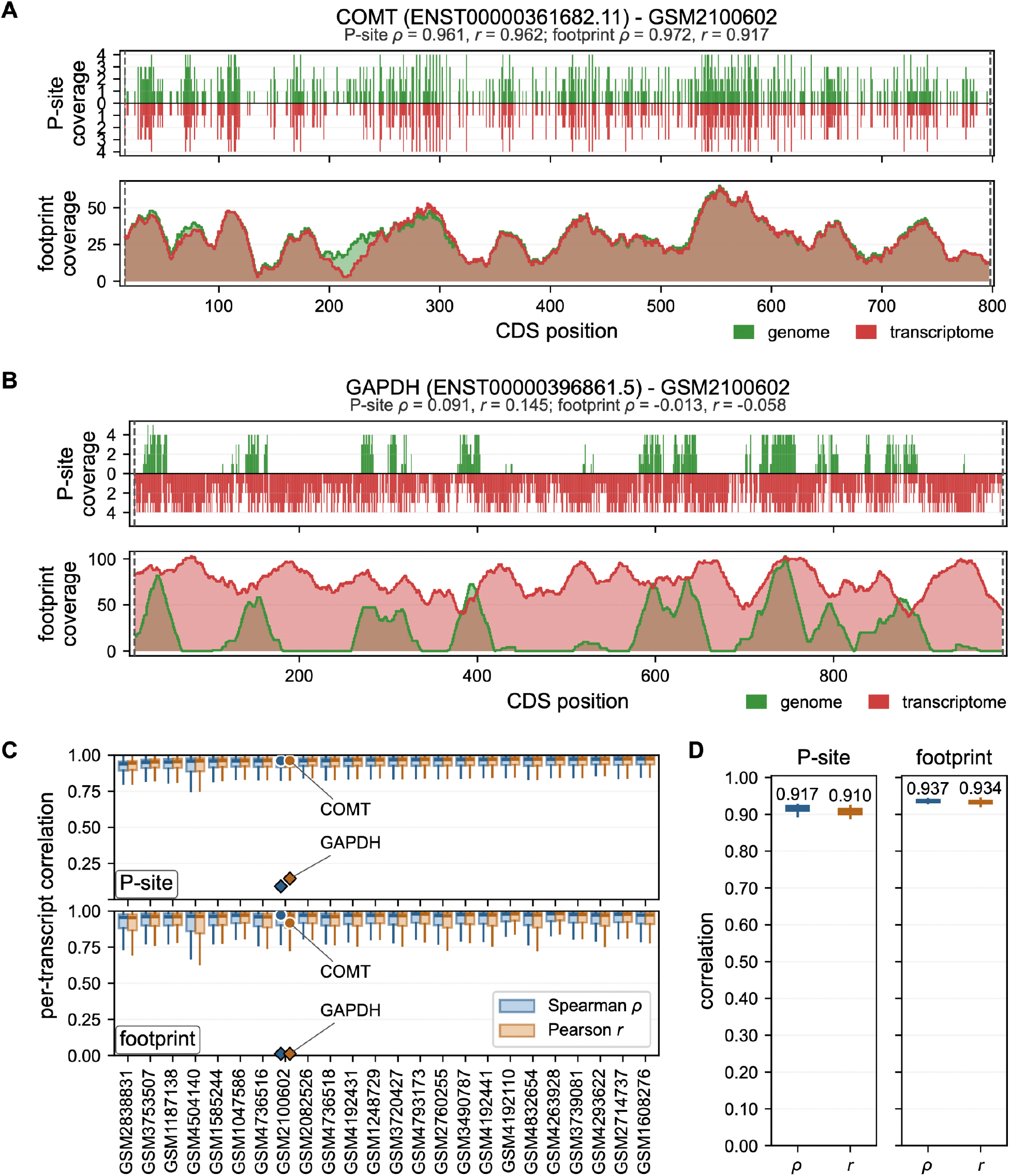
Genome and transcriptome alignments produced concordant nucleotide-level CDS coverage. Genome alignments were restricted to uniquely mapped reads and transcriptome alignments to MAPQ ≥ 42. Coverage was compared at matched CDS positions after excluding 15 nt from each end. Ribosomal P-site coverage assigns each RPF to one nucleotide using its read-length-specific offset. Whole-RPF coverage counts aligned nucleotides spanned by each RPF without P-site offsets applied. Spearman ρ was calculated from raw counts, and Pearson r was calculated from log2(count + 1)-transformed counts. **(A)** COMT and **(B)** GAPDH coverage in GSM2100602. Green and red indicate genome- and transcriptome-aligned coverage, respectively. **(C)** Boxplots of transcript-level correlations for each library. Transcripts without coverage in both routes were excluded, and outliers aren’t shown. Diamonds identify GAPDH, and circles identify COMT. **(D)** Library-level correlations after concatenating matched CDS positions. Values above the distributions indicate medians across the 24 libraries.

### 3.3 TE estimates are highly correlated but systematically shifted between alignment routes

Translation efficiency (TE) was estimated for each alignment route (Methods). Within each sample, genome- and transcriptome-aligned TE estimates were highly correlated across transcripts, with a median Spearman ρ of 0.969 and a median Pearson r of 0.966 (Fig 3A).

**Fig 3.**
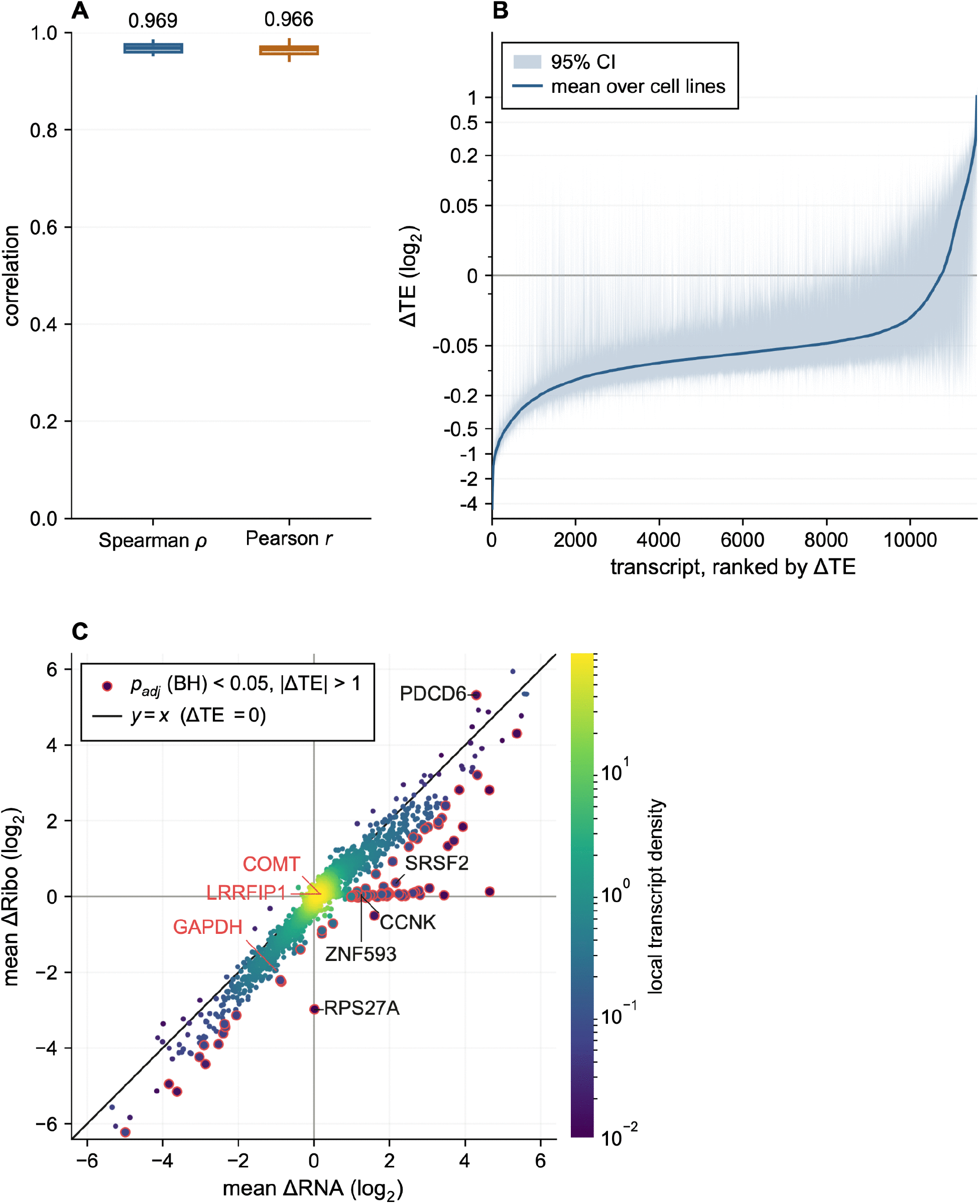
TE estimates are highly correlated but slightly lower for most transcripts with genome alignment. **(A)** Spearman and Pearson correlations between TE estimates obtained from genome and transcriptome alignment across transcripts in each of the 24 cell lines. Values above the distributions indicate medians. **(B)** Mean ΔTE across eligible cell lines for each of the 11,589 transcripts, ranked from lowest to highest. ΔTE was calculated as genome minus transcriptome. Shading indicates transcript-specific 95% confidence intervals. Across transcripts, the median of these means was −0.063 log_2_ units, with an interquartile range of −0.099 to −0.041. The vertical axis uses a symmetric logarithmic scale. **(C)** Mean ΔRNA and mean ΔRibo for each transcript, with both quantities calculated as genome minus transcriptome. The line y = x represents ΔTE = 0. Red outlines identify transcripts with a Benjamini-Hochberg-adjusted p-value below 0.05 and an absolute mean ΔTE greater than 1. Point color indicates local transcript density. GAPDH, COMT, and LRRFIP1 are labeled in red text for reference.

Of the 11,589 transcripts analyzed, 10,727 had a lower mean TE estimate under genome alignment. The median transcript-level mean ΔTE was −0.063 log2 units, with an interquartile range of −0.099 to −0.041 (Fig 3B). We next identified transcripts with consistent differences across samples. Eighty transcripts had an adjusted p-value below 0.05 and an absolute mean ΔTE greater than 1, corresponding to more than a twofold difference in estimated TE between alignment routes (Methods).

To determine whether alignment-associated discrepancies in estimated TE arose from RPF counts, mRNA counts, or both, we compared mean ΔRibo and mean ΔRNA (Fig 3C). Most transcripts had similar assigned counts under both alignment routes. For some transcripts, the alignment route had comparable effects on RPF and mRNA counts, preserving their estimated TE despite variation in the underlying counts. For transcripts with large TE discrepancies, differences in RPF counts primarily drove RPS27A, whereas differences in mRNA counts drove SRSF2, CCNK, and ZNF593. For PDCD6, both count types varied between routes, with a larger effect on RPF counts. Overall, alignment route affected assigned transcript counts even when those effects canceled in the TE ratio, while imbalanced effects between assays produced discrepancies in estimated TE.

### 3.4 Genomic multimapping and transcript representation affect read assignment

To examine sources of alignment-dependent read assignment, we partitioned reads by alignmed source and genomic mapping uniqueness (Methods). Across libraries, a median of 0.72 million more reads aligned to the genome than to the transcriptome (IQR, 0.47 to 0.99 million; Fig 4A). The median proportions of SH-U, SH-M, GO-U, GO-M and TO reads were 75.4%, 9.3%, 9.2%, 4.5% and 1.0%, respectively (Fig 4B). Among SH-M reads, a median of 52.4% had equally scoring genome alignments between a protein-coding gene and a processed pseudogene (IQR, 49.5% to 54.3%; Fig 4C). Across libraries, a median of 28.8% of GO-U reads mapped to exons absent from the selected reference transcript (IQR, 24.8% to 31.0%; Fig 4D).

**Fig 4.**
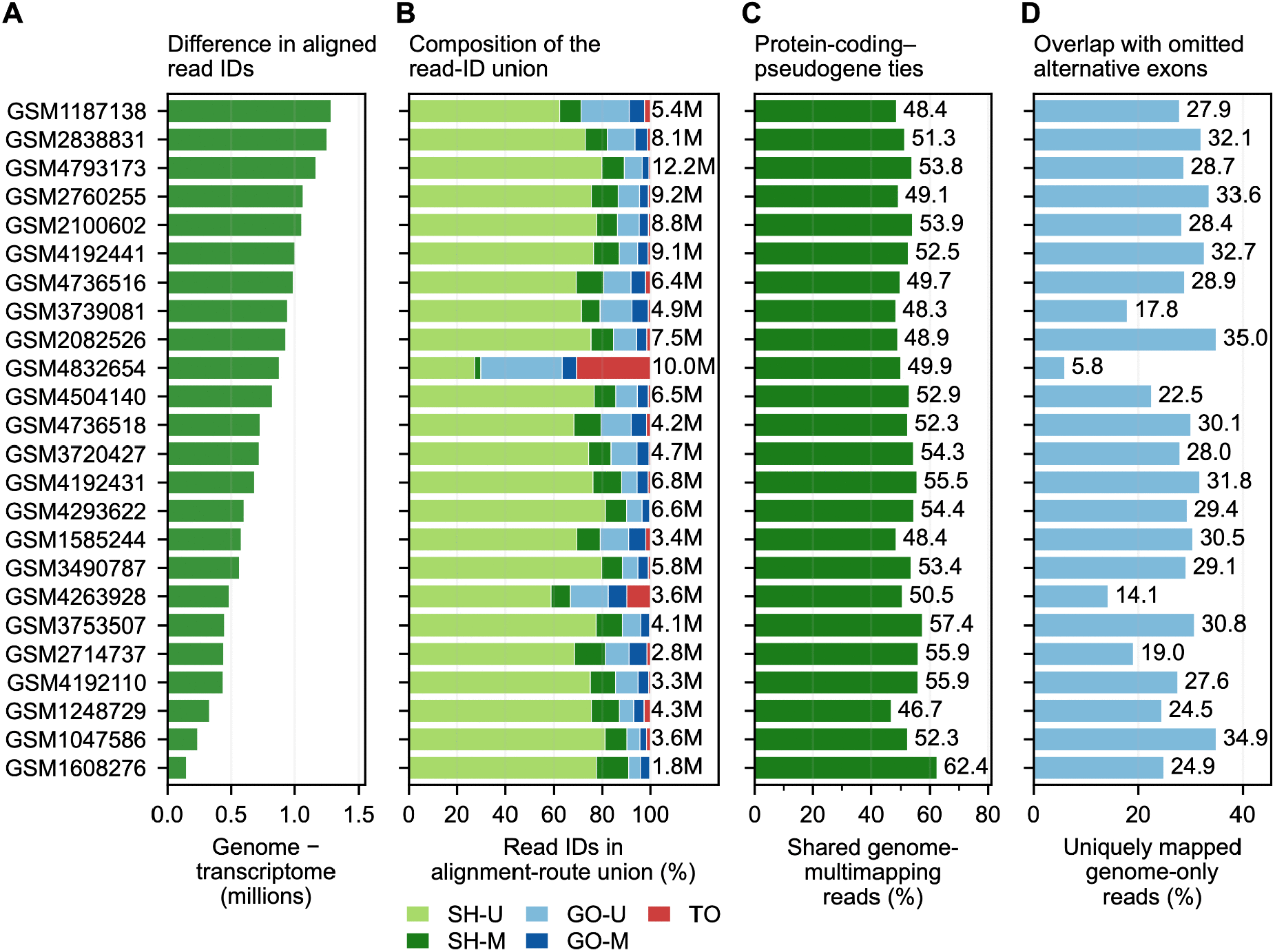
Genomic multimapping and transcript isoform representation contribute to alignment-specific read assignment. **(A)** Difference in the number of distinct mapped RPF read IDs between genome and transcriptome alignment, calculated as genome minus transcriptome. (Median 0.72 million; IQR, 0.47 to 0.99 million). **(B)** Composition of the read-ID union within each library. Categories are shared and genome-unique (SH-U), shared and genome-multimapped (SH-M), genome-only and unique (GO-U), genome-only and multi-mapped (GO-M), and transcriptome-only (TO). Medians were 75.4%, 9.3%, 9.2%, 4.5%, and 1.0%, respectively. Labels indicate the total number of mapped read IDs. **(C)** Percentage of SH-M reads with tied best-scoring alignments between a protein-coding gene and a processed pseudogene. (median 52.4%; IQR, 49.5% to 54.3%). **(D)** Percentage of GO-U reads mapped to exons absent from the selected transcript (median 28.8%; IQR, 24.8%-31.0%).

To illustrate how these patterns varied among genes, we examined COMT and GAPDH in GSM2100602. Of the reads mapped to COMT, 84% were SH-U (Fig 5A), consistent with the coverage profiles observed under the two alignment routes (Fig 2A). Of all read IDs associated with GAPDH under either alignment route, 24% were SH-U and 54% were SH-M reads with equally best-scoring genomic alignments at GAPDH and a processed pseudogene (Fig 5A). These observations help explain why genome coverage and estimated TE were lower for GAPDH when genome analyses were restricted to uniquely mapping reads.

**Fig 5.**
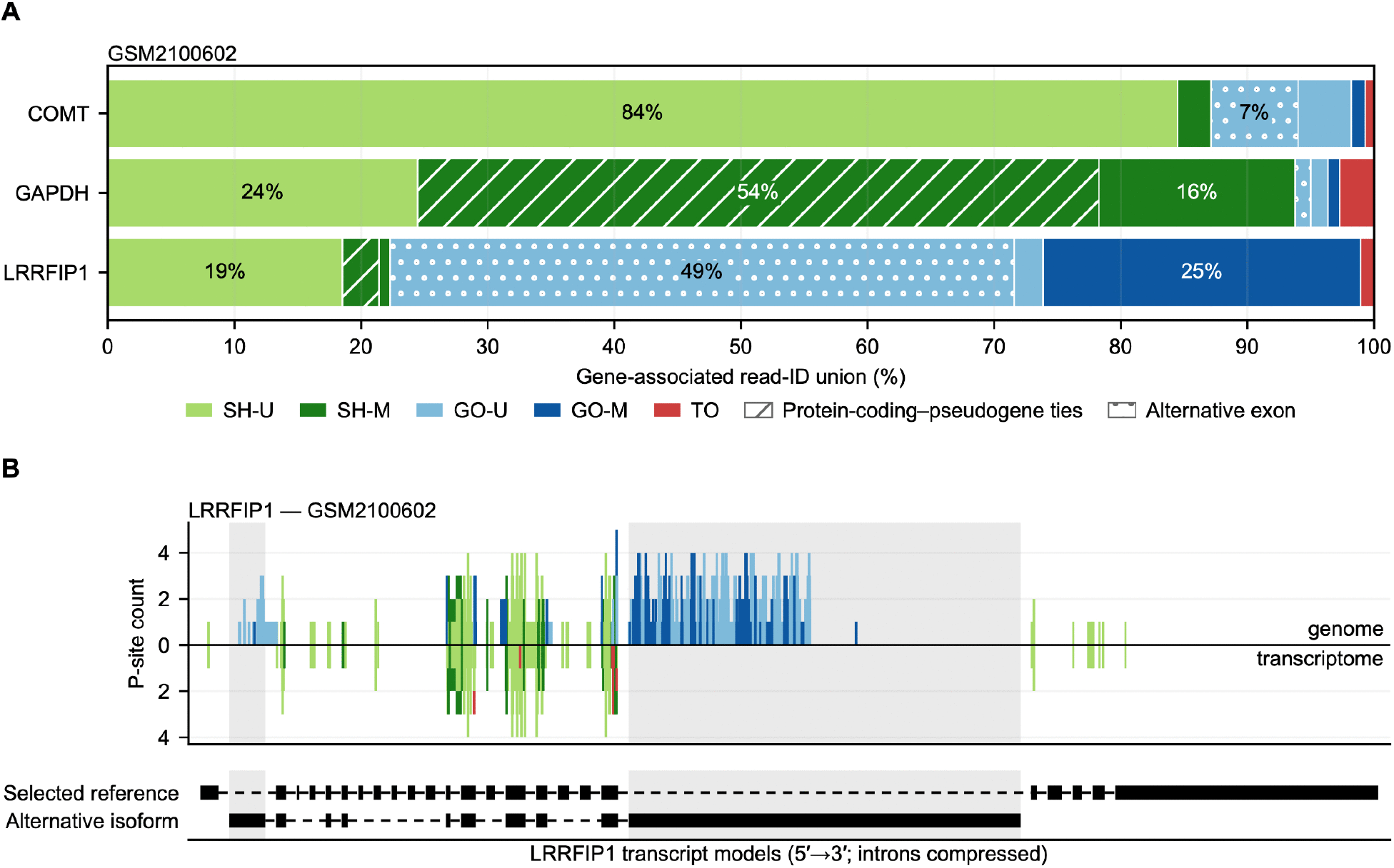
Read assignments at individual genes reflect genomic multimapping and alternative isoforms. Read IDs associated with COMT, GAPDH, or LRRFIP1 in GSM2100602 were classified as shared genome-unique (SH-U), shared genome-multimapped (SH-M), genome-only unique (GO-U), genome-only multimapped (GO-M), or transcriptome-only (TO). **(A)** Percentage of each gene’s distinct read IDs in each category. Diagonal hatching marks SH-M reads with equally best-scoring genomic alignments at the plotted gene and a processed pseudogene. Dotted hatching marks GO-U reads overlapping exonic sequence absent from the selected reference transcript. Selected percentages are labeled. **(B)** Mirrored P-site coverage for LRRFIP1, with genomic coverage above the axis and transcriptome coverage below. Colors correspond to the categories in panel A, without hatching. Coverage used the selected read lengths and inferred P-site offsets for each route. Genomic coverage used unique and multimapped reads with a highest-scoring alignment at LRRFIP1, and transcriptome coverage used alignments with MAPQ ≥ 10. Transcript models are oriented 5′ to 3′ with introns compressed.

LRRFIP1 showed a different pattern related to isoform representation. Among reads assigned to LRRFIP1 by either route, half were GO-U reads in exons absent from the selected reference transcript, and a further 25% were GO-M (Fig 5A). Genome-aligned P-site coverage was observed in exons present in an alternative LRRFIP1 isoform but absent from the selected reference transcript (Fig 5B).

To identify more genes with read-assignment profiles like those of COMT, GAPDH, and LRRFIP1, we hierarchically clustered 11,864 genes in GSM2100602 (Fig 6A). Clusters 1 through 4 contained 9,699, 811, 690, and 664 genes, respectively. COMT, GAPDH, and LRRFIP1 belonged to clusters 1, 2, and 3, respectively (Fig 6B). Read-assignment proportions varied among genes within each cluster (Fig 6C). The mean gene-level proportions of the largest category were 83% SH-U in cluster 1, 68% SH-M in cluster 2, 68% GO-U in cluster 3, and 73% GO-M in cluster 4 (Fig 6D). Predicted pseudogenes were most common among genes in cluster 2 (40%, Fig 6E). In cluster 3, a median of 22% of annotated exonic sequence was absent from the selected reference transcript, similar to the 23%–32% medians in the other clusters (Fig 6F). A median of 65% of selected-transcript exonic sequence in cluster 3 was also present in another gene’s selected transcript, compared with 0% in the other clusters (Fig 6G).

**Fig 6.**
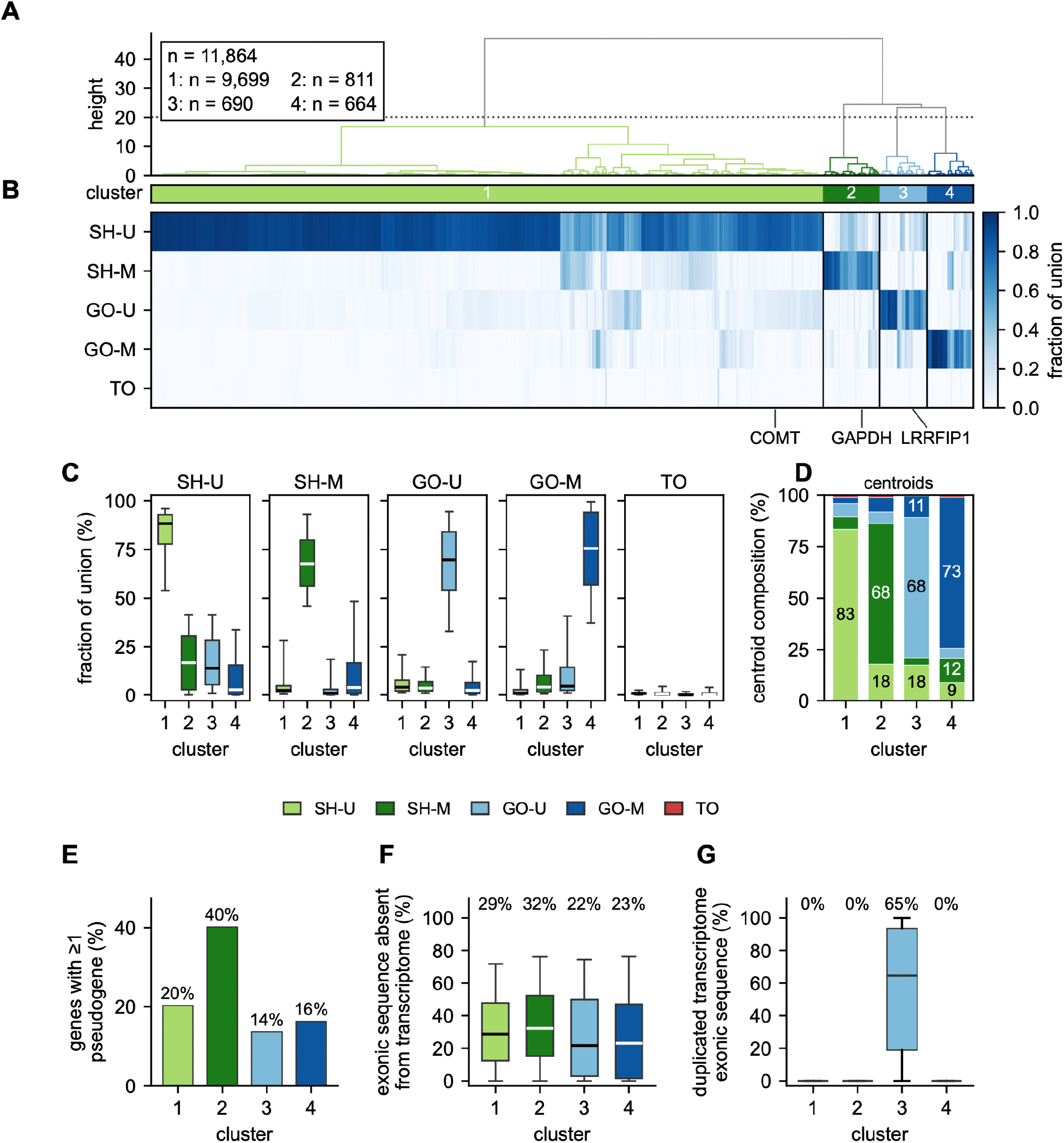
Hierarchical clustering of genes by read-assignment composition. In GSM2100602, 11,864 genes had at least 100 distinct read IDs associated with either alignment route. Read IDs were classified as shared genome-unique (SH-U), shared genome-multimapped (SH-M), genome-only unique (GO-U), genome-only multimapped (GO-M), or transcriptome-only (TO). Transcriptome presence required MAPQ ≥ 10. The proportions of the five categories were clustered using Euclidean distance and Ward’s linkage. **(A)** Dendrogram cut into four clusters. Clusters are colored by the highest mean proportion in each cluster. **(B)** Heatmap of read-assignment proportions for genes, with COMT, GAPDH, and LRRFIP1 labeled. **(C)** Distribution of gene-level category proportions by cluster. **(D)** Cluster centroids shown as mean compositions. **(E)** Percentage of genes in each cluster with at least one predicted pseudogene. **(F)** Percentage of each gene’s annotated exonic sequences absent from the selected reference transcript used for transcriptome alignment. Medians are labeled above boxplots. **(G)** Percentage of each reference transcript’s exonic sequence also present in another gene’s reference transcript on the same strand.

### 3.5 Discussion of Results

For most genes, the choice between genome and transcriptome alignment had little effect on CDS coverage or TE estimates when both routes used stringent mapping-quality filters. However, TE estimates sometimes remained similar because RPF and mRNA counts changed proportionally between alignment routes. Agreement in TE therefore does not necessarily indicate agreement in the underlying abundance estimates. Substantial differences in positional coverage at individual genes also show that the apparent distribution of ribosome occupancy can depend on the alignment route despite broad agreement across transcripts.

The difference in coverage uniformity observed for GAPDH illustrates how alignment choices may alter features used in downstream translation analyses. For GAPDH, excluding reads with equally scoring genomic alignments at processed pseudogenes helps explain the loss of coverage under stringent filtering. This filtering can also alter TE when RPF and RNA counts are affected disproportionately. Although tied alignments alone cannot identify the locus that produced a read, GAPDH is among the most abundant proteins in human cells, whereas its pseudogenes are predominantly processed copies retrotransposed from GAPDH mRNA. These RPFs almost certainly derive from GAPDH, and the transcriptome route assigns them correctly. Unique-only genome alignment discards them, leaving coverage gaps wherever GAPDH sequence is shared with a pseudogene. Translation has been reported for some human pseudogene transcripts [24], but establishing it requires reads that align uniquely to pseudogene-specific sequence, not tied alignments.

LRRFIP1 illustrates another limitation, as similar coverage at matched CDS positions and similar TE estimates exclude additional RPF coverage in alternative exons absent from the selected transcript. This highlights the importance of examining regions beyond selected reference transcripts, particularly when isoform usage varies across tissues and cell types [7,8].

We then extended our individual gene observations by identifying genes sensitive to alignment methodology. As expected, genes with predicted pseudogenes were highly dependent on the reference choice, where exclusion of these competing loci from the transcriptome reference allowed reads with ambiguous genomic alignments to be assigned confidently to selected transcripts. Unexpectedly, the proportion of exons absent from the transcriptome reference did not distinguish the GO-U dominated cluster from others. Instead, genes in this cluster had a greater proportion of exonic sequence shared with other reference transcripts. This reveals a limitation in transcriptome reference, as ambiguity among overlapping transcripts may contribute to the loss of reads during transcriptome alignment filtering. Transcriptome-based ribosome profiling analysis therefore requires careful attention to both transcript inclusion and shared sequences among overlapping and read-through genes, which may reduce alignment confidence. Genomic alignment may help preserve coverage in these regions.

RiboFlow v2 enables assessment of alignment-route sensitivity across coverage, counts, and TE for ribosome profiling experiments. Although we did not fully resolve the source of discrepancies, we identified genes sensitive to alignment and mapping decisions and highlighted how assumptions about transcript representation and multi-mapping affect coverage and abundance estimates. The findings depend on the quality and completeness of the transcript models and genomic annotation used. These requirements may limit comparison in non-model organisms.

## 4. AVAILABILITY AND FUTURE DIRECTIONS

The current release of RiboFlow v2 does not yet store genome-aligned data in the .ribo file format. We plan to implement this extension with a second version of the ribo file format in the near future. The updated format will accommodate genome-aligned data, accompanied by corresponding updates to RiboR, RiboPy, and RiboGraph to support genome-based analyses throughout the Ribo ecosystem.

RiboFlow v2 source code, installation instructions, and example configurations are available at https://github.com/ribosomeprofiling/riboflow and archived on Zenodo [48].

BAM files generated by RiboFlow v2 and used for analysis are deposited on Zenodo at https://doi.org/10.5281/zenodo.22083992 [49].

Analysis scripts to reproduce results, figures, and tables are available on Zenodo at https://doi.org/10.5281/zenodo.23005997 [45].

## Acknowledgments

This work was supported by a National Institutes of Health grant (R35GM150667) and a Welch Foundation grant (F-2027-20260402) (C.C.). An LLM was used to check grammar and suggest clarity improvements in the text. (GPT-5.6 Sol). Claude Code (Opus 5, Anthropic) was used to generate initial drafts of source code, assist with formatting of figure panels, and generate software tests. All outputs were reviewed, edited, and validated by the authors. We thank Hakan Özadam for their feedback and suggestions on the manuscript.

## Supporting Information

**S1.Table RiboBase cell line panel, matched RNA-seq samples, and sample-level quality metrics.**

24 RiboBase samples selected for this study, including GEO accessions for the ribosome profiling and matched RNA-seq libraries, RiboBase quality metrics used for sample selection, selected read-length ranges and periodicity metrics, CDS coverage and read counts across fixed and sample-specific read-length ranges, normalized library depth, and sample annotations.

**Fig S1.Read length selection, P-site offsets, and CDS periodicity agree between the genome and transcriptome alignments.**

24 libraries were aligned to the genome with STAR and to the transcriptome with Bowtie2. Reads that uniquely mapped to the genome and transcriptome alignments with MAPQ ≥ 42 were retained. **(A)** Selected read lengths and inferred ribosomal P-site offsets for each library (rows) and read length (columns). Numbers indicate inferred P-site offsets. **(B)** Difference (genome - transcriptome) in the percentage of P-sites in the first coding-sequence (CDS) reading frame after applying the inferred offsets. Differences are in percentage points. The mean absolute difference was 0.33 percentage points.

